# SAM-DTI: A Spatial Attention Model for Drug-Target Interaction Prediction

**DOI:** 10.1101/2025.08.31.673317

**Authors:** Madhurima Mondal, Shili Wu, Aniruddha Datta

## Abstract

Drug-target protein interaction (DTI) prediction is an important area of research in drug repurposing and discovery. Laboratory experiments for exploring DTI space need a lot of time and money due to the high volume of potential compounds. The past decade’s advancements in deep learning have made progress in efficient prediction of DTIs. This paper proposes a spatial attention model for predicting drug-target interactions (SAM-DTI) from labeled SMILES strings and protein sequences. SAM-DTI (i) tries to encapsulate the input features with more relevance to the biological binding of drug-target interactions, rather than models that solely extract features from individual inputs, and (ii) emphasizes giving dynamic importance to each pair of drug-target interactions. The model follows a knowledge-inspired sub-structure vocabulary to preprocess the inputs. It predicts accurate DTI values from the interactions between each SMILES sub-structure and protein substructure pair. SAM-DTI achieves competitive performance compared to state-of-the-art methods.

## I. Introduction

Despite tremendous improvements in health and biotechnology, drug discovery and development (DDD) still takes expensive financial commitment and time due to the high volume of potential drug compounds [1]. Even though clinical trials are the most reliable part of the DDD process, several opportunities lie in the pre-clinical stage for reducing time and cost [2]. A popular method to measure the therapeutic effects of drug compounds is the drug-target protein interaction (DTI) values [3]. DTIs serve as the fundamental tool in drug repositioning. Scientists must conduct assay experiments to predict DTIs and identify the potential therapeutic compounds accurately. These investigations come with the cost of patients’ safety risk in clinical trials, ethical lapses in xenographic studies, and significant time and expenses in both [1], [4].

Recent developments in deep learning have brought revolutionary technologies to the healthcare domain [5]. One of those technologies is emerging computational models for DTI predictions from vast databases and specific domain knowledge. Since cellular proteins selectively interact with drug molecules to combat diseases [6], DTI prediction becomes a binary classification task for deep learning - determining whether a given pair of drugs and proteins will interact [7]. Deep learning models for DTI prediction involve taking drugs and target proteins as inputs and analyzing their features to determine interaction. Later, these trained models can predict the DTI values of unseen data [1], [8]. In addition to the deep learning techniques such as deep neural networks (DNN) [9], convolutional neural networks (CNN) [10], graph neural networks (GNN) [11], other methods such as recurrent neural network (RNN) [12], [13], long-short term memory (LSTM) [14], [15], and transformer networks [16] have shown significant promise in DTI prediction.

### Recent Work

There are already many impressive and innovative methods for DTI prediction. Early on, similarity score [17], [18] or distance-based [19] algorithms were among the most popular methods for DTI prediction. However, such methods often fail to generalize on unseen data [3]. LASSO- DNN [20] which leverages linear feature selection for DTI prediction faces challenges due to oversimplification. DeepDTI [21], a deep learning-based DTI prediction method uses deep belief networks (DBN) [22] to forecast new DTIs from raw input vectors of drugs and proteins, although it suffers from a limited contextual understanding of sequences. Transformer-based methods like TransDTI [23], and MolTrans [1] solve the previous challenge of context. However, they highly focus on multilayer encoding of individual drug and protein embeddings. Other popular methods like, DeepWalk [24], and GraphDTA [11] that involve the graph neural networks suffer from scalability issues. However, they also face a struggle to capture the complexity and fully address several drawbacks:

- **Lack of biological binding mimicry:** Popular drugtarget interaction (DTI) prediction methods, such as DeepDTA [25], WideDTA [26], and DeepDTI [21] apply deep learning tools to learn drug and protein features but often fail to accurately simulate the biological reality of drug-target binding. These neural networks usually concentrate on sequence-based feature extractions and ignore the spatial relationship between the drug and protein vectors. As a result, they often fail to recognize the key binding features that affect the interactions the most. In cellular systems, drug-protein bindings are highly localized and typically occur at key binding pockets of proteins, i.e. only a few regions are responsible for the interaction. Once the drug binds to these key regions, it initiates a sequence of downstream molecular events that ultimately lead to combating diseases [27], [28]. The models above often overlook this fact and miss finer features in DTI prediction. These limitations often make the above models less practical and unreliable for real world applications.
- **Potential for Information Loss in Early Processing:** Many existing models such as MolTrans [1], DeepChem over emphasize deep encoding techniques (e.g. transformers or RNN) of drug and protein sequences before considering their interaction. These models do not consider vital interaction patterns that can be lost during the processing of drug and protein sequences in the early stage. Feeding the drug and protein sequence through multiple layers independently carries the risk of over-abstracting the individual features. Transformers are particularly good at modeling long-range dependencies inside sequences, as noted in [16] and [30], but are less effective at representing cross-sequence interactions when sequences are processed independently.

### Our Contribution

To address the aforementioned challenges, we propose a spatial attention-based method that is applied to the initial interaction map of drugs and proteins to predict DTIs. This drug-protein binding-inspired method utilizes pattern-based sub-structure tokenization, ESPF [31] of drug and protein sequences. After tokenization, the method forms the drug-protein interaction matrix which is then analyzed with spatial attention to capture local and global features. Our method, **S**patial **A**ttention **M**odel for **DTI** prediction (SAM-DTI) solves the above challenges in the following manner:

- **Biological binding for accurately simulating drugtarget interaction:** Biologically, applying features in the drug-protein interaction matrix would be the correct approximation for DTI prediction. The attention mechanism emphasizes cross-interactions between drugs and proteins that are biologically more important. The twodimensional spatial kernel allows complementarity and proximity between certain drug and protein substructures that bind to each other.
- **Preserving critical interaction information through early integration:** In a deep learning setup the early layers extract the low-level features and the later layer extracts the high-level features. Our technique maintains necessary spatial and functional relationships early on, generating the interaction map right after the embeddings are generated. This helps the model to preserve relevant low-level interacting features.
- **Adaptive and Context-Sensitive:** Unlike CNNs with their fixed convolutional filters, spatial attention is context-sensitive and adaptive, which means it can change its emphasis depending on the particular drug-protein pair being studied. Because of its adaptiveness, the model can identify different levels of interaction, depending on the distinct structural and chemical characteristics of the molecules, which enhances the precision of binding predictions made for a range of drug-protein interactions.

The paper is organized as follows. In Section II, we discuss representing drug-target protein interaction as a binary classification problem. Then, in Section III, we present our materials and proposed methods. The results of several experiments we performed are included in Section IV. Finally, Section V concludes the paper by summarizing the main results and outlining the topics for future research.

## II. ProblemDefinition

DTI can be formulated as a binary classification problem whose goal is to predict whether a given drug interacts with a specific protein. Suppose, *D* denotes the space of SMILES (Simplified Molecular Input Line Entry System) sequences of drugs and *P* denotes the space of amino acid sequences of proteins.

### A. Input Representation

#### Drug Representation

A drug molecule *d* ∈ *D* is represented by a sequence of sub-structures derived from its SMILES string:

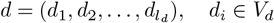

where *l_d_* is the length of the drug sequence and *V_d_* is the vocabulary of possible sub-structures.

#### Protein Representation

A protein *p* ∈ *P* is represented by a sequence of amino acids:

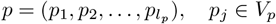

where *l_p_* is the length of the protein sequence and *V_p_* is the vocabulary of possible amino acid sequences.

### B. Output function

The objective of the DTI task is to predict whether a drug *d* and a protein *p* interact. This interaction can be represented as a binary mapping *f* (*d, p*): *y* = *f* (*d, p*) ∈ {0, 1} where *y* = 1 indicates that the drug interacts with the protein, and *y* = 0 indicates no interaction.

## III. Materials andMethods

### A. Sequence processing

We utilized Explainable Substructure Partition Fingerprint (ESPF) [31] to tokenize input sequences. ESPF is inspired by subword tokenization [32] in natural language processing and derives tokens by using byte pair encoding (BPE) [33]. This has proved to capture meaningful substructures by iteratively merging the most frequent pairs of tokens in a large set of sequences [1], [34]. In this paper ESPF breaks down the drugs and protein sequences in relevant substructures and encodes them to overcome the drawbacks of traditional molecular fingerprints [31].

### B. SAM-DTI model

We propose a Spatial Attention Model for DTI prediction that consists of an embedding layer, followed by an interaction map. Unlike conventional models that process sequences independently, SPTM-DTI, shown in Fig. 1 focuses on crucial regions of drug-protein binding with the application of spatial attention to the interaction matrix and helps in mimicking realworld molecular interactions where certain substructures are concerned with the actual binding process. It also addresses the problem of late-stage integration by creating an interaction map upfront and maintaining the necessary interaction information throughout the process.

**Fig. 1.**
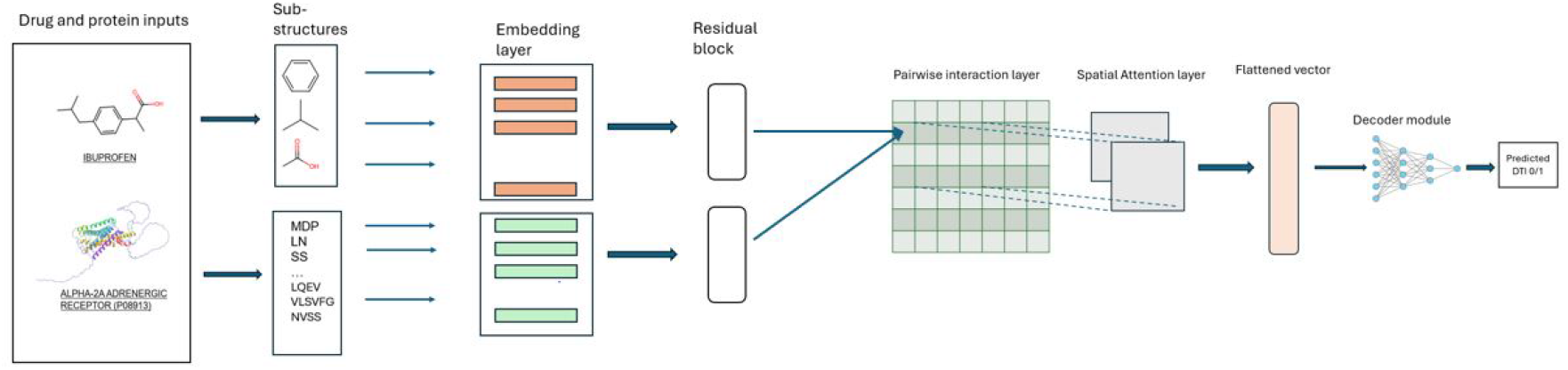
This is the proposed SAM-DTI architecture. The inputs to this architecture are SMILES strings of drugs and amino acid sequences of proteins. We represented the pair-wise interaction matrix in the green-grided cell.

#### 1) Embedding

Each drug and protein sequence is first embedded into a latent space. Given a drug SMILES string and a protein amino acid sequence, we use a token dictionary to encode the tokens. The drug sequence *d* of length *l_d_*, and a protein sequence *p* of length *l_p_* are embedded in *E_d_* and *E_p_* where 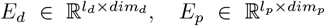 where *dim_d_* and *dim_p_* are the embedding dimensions of the drug and protein, respectively. Embeddings vectors are obtained from a learnable matrix 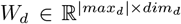 for drugs and 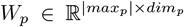 for proteins, where *max_d_* and *max_p_* represent embedding dimension of drug and protein sequences respectively. The embedding for each sub-structure can be formalized as: 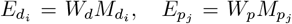 where 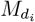 and 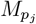 are the one-hot encoded representations of the *i*-th drug and *j*-th protein sub-structures.

##### Positional embedding

We include the positional embeddings *P_d_* and *P_p_* into the model to capture the sequential nature of drugs and proteins because the order of sub-structures is critical in determining the overall properties of the molecule. We compute the positional embedding as: 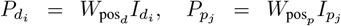 where 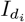 and 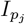 represent the positional indices of the drug and protein substructures. The sum of the content and positional embeddings yields the final embeddings for both sequences.:

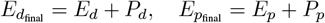

##### Pairwise embedding matrix formation

We construct a pairwise interaction matrix from the outer dot product of drug and protein embeddings to model the interactions between drug and protein sub-structures. This matrix represents the pairwise interaction of the drug and protein tokens:

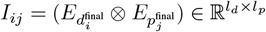

where ⊗ denotes the outer dot product, measuring the interaction between the *i*-th sub-structure of the drug and the *j*-th sub-structure of the protein.

##### 2) Attention block

#### Spatial Attention

The drug-protein interactions include local relationships such as adjacent atoms in a molecule, and global relationships like distant substructures contributing to a binding pocket. To capture these interactions, we apply a spatial attention mechanism to the pairwise interaction matrix *I*, which focuses on both local and global features in the interaction map. Our spatial attention mechanism starts by applying convolutional layers on the interaction matrix *I* to generate the query, key, and value tensors. The convolution layer allows the model to capture **low-level spatial dependencies** by focusing on nearby regions in the interaction matrix. Specifically, the query *Q*, key *K*, and value *V* tensors are computed as follows:

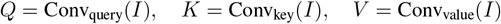

where Conv_query_, Conv_key_, Conv_value_ are 2D convolutional operations with learnable filters. The convolution kernel slides over the spatial dimensions of the interaction map *I*, allowing the model to capture local patterns by extracting spatial features from small patches of the matrix. After generating the query, key, and value tensors, the spatial attention mechanism proceeds to compute the attention score for each sub-structure pair:

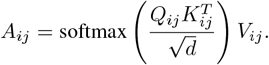

Here, *A_ij_* is the attention score for the interaction between sub-structures *i* and *j*, and *d* is the dimensionality of the key. The softmax function ensures that the attention scores are normalized across the interaction matrix. The attention mechanism, combined with the initial convolutional layers, enables the model to dynamically assign importance to specific regions in the interaction map, thus capturing both **local interactions** (within neighboring sub-structures) and **global interactions** (between distant sub-structures). The final output of the spatial attention mechanism is a refined interaction map that integrates both local and global information, providing a more comprehensive representation of drug-protein interactions.

##### Self-Attention and residual connections

To improve the model we used self-attention [16], [35], which allows each sub-structure to attend to all other sub-structures within the same sequence. This enables the model to learn long-range dependencies. Mathematically, the self-attention operation is defined as:

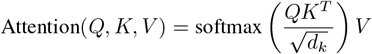

where 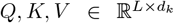 are the query, key, and value matrices, and *d_k_* is the dimensionality of the key. This attention mechanism is followed by residual connections [36], which ensure the stability of gradient flow by adding the input directly to the output of each self-attention block:

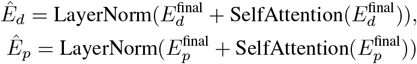

This operation preserves the original embedding while augmenting it with global contextual information.

##### Interaction prediction with MLP

Once the interaction matrix is refined using spatial attention, it is flattened into a vector and passed through a multi-layer perceptron (MLP) to predict the likelihood of a drug-target interaction:

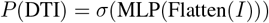

where *σ* is the sigmoid activation function, and the MLP consists of several fully connected layers with ReLU activations and dropout regularization to prevent overfitting. The output of the MLP is a scalar value representing the probability of interaction. We utilize a weighted binary cross entropy loss for our model to mitigate the effects of the unbalanced data.

### C. Implementation

The SAM-DTI is implemented using PyTorch [37]. Drug and protein sequences are tokenized using the ESPF tokens mentioned in [1]. The embedding size is set to 128 for both, with a maximum sequence length of 50. Multi-head self-attention of eight heads is applied to both drug and protein embeddings. The spatial attention key captures binding features from the interaction matrix using convolutional layers. The Adam optimizer with a learning rate of 0.0005121 is used for 100 epochs with a batch size of 64. We applied a dropout of 0.37 to prevent overfitting. We tuned the hyperaparameters using Optuna [38].

## IV. Results

### A. Experimental set up

#### Dataset

We performed our experiments on the DAVIS dataset [39] which has 68 drug nodes and 379 protein targets. The dataset contains *K_d_* values for 25,772 DTI pairs. The *K_d_* values are binarized with a threshold of 30. DAVIS data is highly unbalanced with positive and negative samples. Hence we collected negative sampled [40] data from [1]. The balanced data had 1043 positive and 1043 negative samples in the training data, 160 positive and 2846 negative samples in the validation data, and 303 positive and 5708 negative samples in the test data. We implemented our model on other two datasets, BindingDB [41] and BIOSNAP [42]. The dataset statistics are given in Table I.

**TABLE I.**
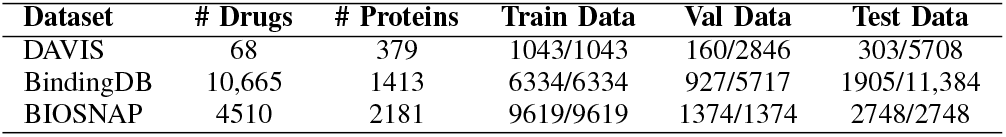
Dataset statistics.

#### Metrices and Evaluation strategies

We use ROC-AUC (Area Under the Receiver Operating Characteristic Curve) and PR-AUC (Area Under the Precision-Recall Curve) to check the performance of the model of binary classification. Traditionally, the performance of models is evaluated using ROC-AUC, where values closer to 1 indicate better performance. In particular, this is even more informative if the classes are balanced. PR-AUC is more relevant to the performance concerning the positive class and thus, by nature, best suited for the unbalanced datasets. A higher PR-AUC value is indicative of better precision-recall trade-offs. We further report the sensitivity or recall, specificity, and F1 score to comprehensively evaluate this model’s performance. Each experiment is repeated five times, considering five different random states for robustness.

### B. Baselines

We compared the performance of our model with the following baseline methods:

- **LR:** Logistic regression [43] was applied on concatenated drug and protein vectors processed using ESPF and run over 1000 iterations.
- **DNN:** ESPF processed the drug and protein sequences were concatenated, and passed through a fully connected DNN with two hidden layers. The DNN applied ReLU activation and dropout to prevent overfitting.
- **DeepPurpose:** [44] We used the DeepPurpose library to load and preprocess the datasets, applying MPNN [45] for drug encoding and CNN for protein-encoding. The model was trained and evaluated for drug-target interaction prediction.
- **DeepConvDTI:** [10] We implemented this model using Keras with convolutional layers for protein sequences and dense layers for drug features. The model includes batch normalization, dropout, and fully connected layers for interaction prediction.
- **MolTrans:** [1] We implemented this model for 5 random runs with hyperparameters mentioned in the paper. The model was trained over ESPF vocabulary.

### C. SAM-DTI achieves competitive performance

Our model shows improved performance on the DAVIS dataset. SAM-DTI outperforms various baseline models in several metrics such as AUC and precision-recall and we summarize the comparisons in Table II. Interestingly we found a higher PR-AUC and sensitivity compared to others, reflecting the capability of the model for selecting true positives over false positives more efficiently. In other words, it identifies true interactions more precisely from actual positives, which is useful for drug repurposing or other tasks in biological data when the finding of true interactions is important. We represent it as a highly effective method for drug-target interaction prediction.

**TABLE II.**
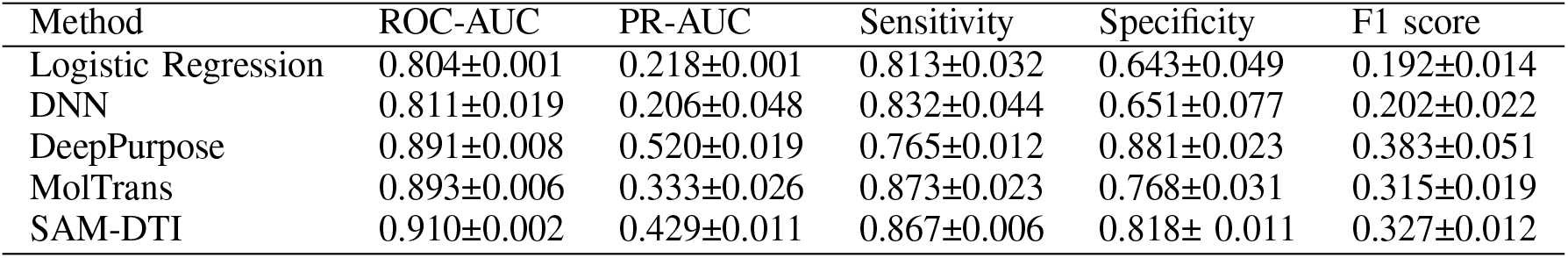
Performance comparison(five random runs)

| Method | ROC-AUC | PR-AUC | Sensitivity | Specificity | F1 score |
| --- | --- | --- | --- | --- | --- |
| Logistic Regression | 0.804±0.001 | 0.218±0.001 | 0.813±0.032 | 0.643±0.049 | 0.192±0.014 |
| DNN | 0.811±0.019 | 0.206±0.048 | 0.832±0.044 | 0.651±0.077 | 0.202±0.022 |
| DeepPurpose | 0.891±0.008 | 0.520±0.019 | 0.765±0.012 | 0.881±0.023 | 0.383±0.051 |
| MolTrans | 0.893±0.006 | 0.333±0.026 | 0.873±0.023 | 0.768±0.031 | 0.315±0.019 |
| SAM-DTI | 0.910±0.002 | 0.429±0.011 | 0.867±0.006 | 0.818± 0.011 | 0.327±0.012 |

### D. SAM-DTI’s performance over different datasets

We performed our experiment over three datasets DAVIS, BindingDB, and BIOSNAP. We summarize our findings in Table III. The results showed our method shows promising performance to different datasets with different data balances.

**TABLE III.** Performance comparison(five random runs)

| Dataset | ROC-AUC | PR-AUC | Sensitivity | Specificity | F1 score |
| --- | --- | --- | --- | --- | --- |
| BIOSNAP | 0.890± 0.003 | 0.900±0.002 | 0.799±0.008 | 0.841± 0.008 | 0.817±0.004 |
| DAVIS | 0.910±0.002 | 0.429±0.011 | 0.867±0.006 | 0.818± 0.011 | 0.327±0.012 |
| BindingDB | 0.917±0.002 | 0.646±0.011 | 0.908±0.008 | 0.785± 0.009 | 0.569±0.006 |

### E. Understanding sub-structure interactions

We paid attention to the pairwise interactions between drug and protein substructures which allows the model to detect significant interaction pairs. We investigated the attention scores on different interactions. More precisely, we showcase a heatmap of interactions among the first 5 drugs and 10 protein substructures of a certain drug-protein pair in Fig. 2. The darker blocks denote stronger and more critical interactions in the prediction of DTI values. For example, drug substructure “CCN2C” and protein substructure “PF,” indicate more influential binding than substructures “=CC=CC” and “PF”. Other significant pairs include “CCN2C” with “KIL”, “=C(C” with “FMG, and “=C(C” with “QQ”. They indicate relatively stronger connections in DTI predictions for the corresponding pairs. In contrast, lighter regions such as “CN(C)CC1” with “LLE” or “CCN2C” with “W”, suggest weaker and less influential interactions. Interestingly, interaction pairs such as “CCN2C” with “YY”, “=C(C” with “YY”, or “=C(C” with “KIL” with white regions, hint at almost negligible attention score. Since we perform an attention mechanism on the interaction matrix, picking up interaction pairs allows our model to detect more biologically relevant interactions and filter out less important ones.

**Fig. 2.**
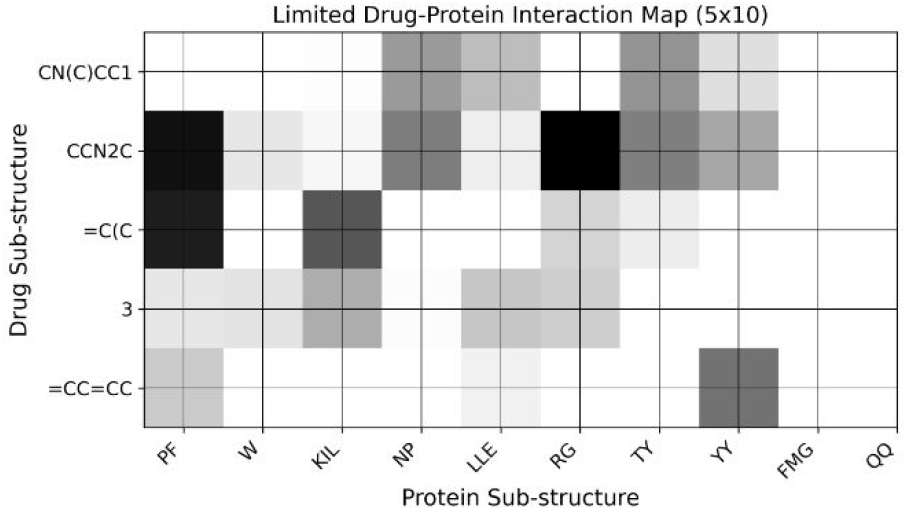
Visualization of spatial attention output on a 5X10 grid

### F. Drug Repurposing

Our drug-target interaction prediction model is also useful in drug repurposing by identifying new interactions between existing drugs and target proteins. For example, we applied our model to curcumin [46], a natural compound, to explore potential new uses. First, we tokenized Curcumin’s SMILES representation and encoded it to make it eligible for feeding in the model. Next, we filtered all the unique proteins from our DAVIS test dataset. We added a sigmoid function to the model output to predict the probabilities between 0 and 1, of curcumin’s interactions with the proteins. This allows us to assess how likely curcumin is to interact with proteins linked to diseases. We considered the five highest probabilities to select the proteins with the strongest interactions. We tracked the gene symbols of proteins and requested the proteins’ UniProt IDs [47] by calling UniProt API. We performed a similar experiment on EGCG and Melatonin. We summarize the list of proteins and diseases associated with Curcumin, along with EGCG and Melatonin in Table IV.

**TABLE IV.** RepurposedDrugs, TargetedProteins, AND RelevantDiseases.

| Repurposed Drug | Targeted Proteins | Relevant Diseases |
| --- | --- | --- |
| Curcumin | AURKC, ABL1<br>FLT4, DDR1 | Lymphatic disorder,<br>Breast cancer, Fibrosis |
| EGCG | DDR1, GSK3A | Breast Cancer, Fibrosis,<br>Neurodegenerative Diseases |
| Melatonin | PRKCE, SLK | Breast Cancer, Prostate cancer,<br>Neurological disorder |

### G. Abalation study

We performed an ablation study to identify important parts of the model. Removal of individual parts affected the ROC-AUC a little. This suggests that the proposed spatial attention approach is sufficient for predicting high ROC-AUC cases. We summarize the findings in Table V.

**TABLE V.**
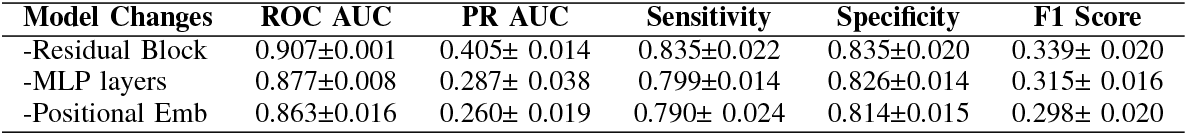
AblationStudyResults forVariousModelChanges.

| Model Changes | ROC AUC | PR AUC | Sensitivity | Specificity | F1 Score |
| --- | --- | --- | --- | --- | --- |
| -Residual Block | 0.907±0.001 | 0.405± 0.014 | 0.835±0.022 | 0.835±0.020 | 0.339± 0.020 |
| -MLP layers | 0.877±0.008 | 0.287± 0.038 | 0.799±0.014 | 0.826±0.014 | 0.315± 0.016 |
| -Positional Emb | 0.863±0.016 | 0.260± 0.019 | 0.790± 0.024 | 0.814±0.015 | 0.298± 0.020 |

#### Removal of decoder layer

We first removed the decoder layer. Surprisingly, the model showed improved ROC-AUC but suffered from instability in the loss curve. Then we experimented by removing different layers of the decoder. We found, that removing two layers from the decoder reduced the model’s performance.

#### Removing the Residual Block

Next, we discarded the residual block after the embedding layer for the ablation study. We found the ROC-AUC remained nearly the same, but the drop in sensitivity suggests that residual connections help capture more relevant features. Specificity and F1 Score also saw slight changes. This highlights that the residual block plays an important role in improving the overall model balance.

#### No Positional Embedding

Our embedding layer contained contextual and positional embedding. Removal of the positional embeddings decreased the ROC-AUC by 1%, suggesting that positional embeddings did not play a very important role in ROC-AUC. Surprisingly, sensitivity decreased the most in this experiment. This indicates that positional embeddings help the model in identifying relevant interactions more effectively.

## V. Conclusion

This paper introduced the Spatial Attention Model for Drug-Target protein Interaction prediction, nicknamed SAMDTI. Our model appears to be a novel approach to drug repurposing and discovery. Our model combines knowledge-inspired tokenization with deep learning techniques. SAM-DTI efficiently captures crucial interactions between drug and protein sub-structures, outperforming the existing state-ofthe-art methods. The key point of our study was to focus on biologically relevant aspects of DTI predictions.

Our findings reveal that the SAM-DTI not only improves prediction accuracy but also provides deeper insights into the mechanism of drug-protein interactions that conventional models usually neglect. Applying spatial attention to the interaction matrix of drugs and proteins, SAM-DTI retains vital interaction information, thereby enhancing the reliability and practical utility of the predictions. This approach also allows for adaptive and context-sensitive predictions that cater to the unique structural and chemical properties of each drug-protein pair.

## VI. BIOGRAPHY SECTION

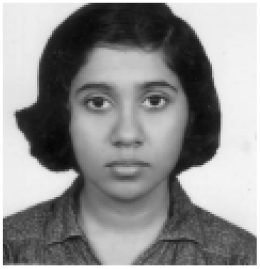

**Madhurima Mondal** is a Ph.D. student in the Genomic Signal Processing Lab at the Department of Electrical and Computer Engineering at Texas A&M University, College Station, USA. She did her bachelor’s and master’s in Electronics and Electrical Communication Engineering from the Indian Institute of Technology, Kharagpur in India. She has been a research intern at the University of Illinois at Urbana Champaign, USA, and a deep learning intern at Samsung R&D Institute, India. She is a recipient of the Indian National Talent Search Examination (NTSE) and KVPY awards. She stood in position 2 (1st among girls) in the Regional Mathematics Olympiad (RMO), India conducted by Homi Bhabha Centre for Science Education (HBCSE).

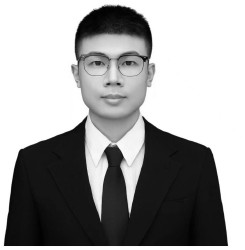

**Shili Wu** is a Ph.D. student in the Genomic Signal Processing Lab at the Department of Electrical and Computer Engineering at Texas A&M University, College Station, USA. He received his B.S. degree from Case Western Reserved University and M.S. degree from Columbia University. His research Interests includes reinforcement learning and control application. He is also interested in applying the machine learning techniques into biography. He has authored several papers in these areas and is an IEEE member.

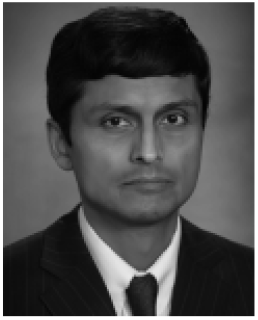

**Aniruddha Datta (Fellow, IEEE)** received the BTech degree in electrical engineering from the Indian Institute of Technology, Kharagpur, India, in 1985, the MSEE degree from Southern Illinois University, Carbondale, Illinois, in 1987, and the MS (applied mathematics) and PhD degrees from the University of Southern California, Los Angeles, California, in 1991. In August 1991, he joined the Department of Electrical and Computer Engineering, Texas A&M University where he is currently the J. W. Runyon, Jr. ‘35 professor II and Co-Director of Graduate Programs. His research interests include adaptive control, robust control, PID control, and Genomic signal processing. He has authored or coauthored five books and more than 200 journal and conference papers on these topics. He has served as an associate editor of the IEEE Transactions on Automatic Control (2001-2003), the IEEE Transactions on Systems, Man and Cybernetics-Part B (2005-2006), the IEEE Transactions on Biomedical Engineering (2013-2015), the EURASIP Journal on Bioinformatics and Systems Biology (2007-2016), the IEEE Journal of Biomedical and Health Informatics (2014-2016), the IEEE/ACM Transactions on Computational Biology and Bioinformatics (2014-2017), the IEEE Access (2013-2022) and is currently serving as an Academic Editor for PLOS One.

